# Solute exchange through gap junctions lessens the adverse effects of inactivating mutations in metabolite-handling genes

**DOI:** 10.1101/2022.03.15.484462

**Authors:** Stefania Monterisi, Johanna Michl, Amaryllis E. Hill, Alzbeta Hulikova, Gulnar Abdullayeva, Walter F. Bodmer, Pawel Swietach

## Abstract

Experimental inactivation of certain genes involved in metabolism attenuates cancer cell growth *in vitro*. However, loss-of-function mutations in metabolic pathways are not negatively selected in human cancers, indicating that these genes are not essential *in vivo*. We hypothesize that spontaneous mutations affecting metabolic pathways do not necessarily result in a functional defect because affected cells may be rescued by exchanging metabolites with neighboring wild-type cells via gap junctions. Using fluorescent substances to probe inter-cellular diffusion, we show that colorectal cancer (CRC) cells are coupled by gap junctions assembled from connexins, particularly the constitutively expressed Cx26. In co-cultures of wild-type cells with cells that had inactivated components of pH regulation (*SLC9A1*), glycolysis (*ALDOA*), or mitochondrial metabolism (*NDUFS1*), we show that diffusive coupling was able to rescue the functional defect associated with the inactivation of metabolite-handling genes. Function rescue was dependent on Cx26 channels and reduced phenotypic heterogeneity among cells. Since the phenotypic landscape did not map onto genotype, an individual cell should not be considered as the unit under selection, at least in the case of metabolite-handling processes. Our findings can explain why certain loss-of-function mutations in genes, previously ascribed as being ‘essential’, do not influence the growth of human cancers.

## INTRODUCTION

The role of somatic evolution in advancing cancers is well-established (1, 2) and can be modelled mathematically (3, 4). Its central paradigm asserts that mutations benefiting cancer cells undergo positive selection and appear enriched in cancers. Cells carrying such genetic changes may evade normal checks and controls that restrict proliferation (5), or become resistant to harsh microenvironments (6). Conversely, mutations that inactivate critically important processes are predicted to emerge less frequently than the random mutation rate (7). *In vitro* knockout screens have identified genes labeled as essential for cell survival (8, 9), for which inactivating mutations are predicted to undergo negative selection in cancers (10, 11). These genes include those involved in metabolism (12, 13). The notion of ‘essential genes’ has gained much traction, despite the scarcity of negative selection *in vivo*, equating to only tens of genes (14) such as those involved in protein synthesis and immune-exposed epitopes, but not metabolite handling (10). The reason for the discrepancy between the interpretation of *in vitro* screens and *in vivo* observations remains unclear.

We hypothesize that mutations leading to the inactivation of key metabolic pathways do not incur functional deficits in mutated cells because diffusive exchange with neighboring wild-type cells provides access to operational proteins. Many tissues are coupled by connexin-assembled gap junctions that allow metabolite exchange across the continuous cytoplasmic space (15–17). Thus, a spontaneous loss-of-function mutation in a cell within a coupled network will not be deleterious when the affected cell is able to access metabolite-handling enzymes or transporters in neighboring cells. If, however, the same mutation is introduced experimentally in all cells, as is common with *in vitro* assays, diffusive exchange cannot restore metabolic function, irrespective of coupling strength. We speculate that the diffusive exchange of metabolites via gap junctions explains why some metabolic processes, deemed to be essential for survival *in vitro*, do not undergo negative selection in cancers. When a mutation-carrying cell resorts to neighboring cells for functional rescue, the emergent phenotype will not align with genotype. This scenario contradicts a key assumption of tumorigenesis models, that mutations act in a cell-autonomous fashion. Exceptions to this paradigm have been postulated for autocrine interactions (18), but the role of gap junctional connectivity has not been explored. We argue that for certain processes, like those handling diffusible metabolites, it is not appropriate to consider a cell as the unit under selection (19).

This study used colorectal cancer (CRC) cells to test our hypothesis that diffusive exchange can rescue cells carrying inactivating mutations in apparently critical genes. We first assessed the degree of cell-to-cell coupling in a panel of CRC cells and identified the major connexin isoforms responsible for assembling these conduits. We then genetically inactivated metabolic processes that play important roles in cancer, to test whether coupling onto wild-type cells can compensate the functional deficit. The inactivated functions included *(i)* Na^+^/H^+^ exchanger-1 (*SLC9A1*), a pH regulator at the plasma membrane that is critical for survival under acidic conditions (20); *(ii)* aldolase A (*ALDOA*), a glycolytic enzyme that is critical for the Warburg effect (21); *(iii)* NADH:ubiquinone oxidoreductase core subunit S1 (*NDUFS1*), part of complex I required for oxidative phosphorylation (22). In support of our hypothesis, we found that co-culturing genetically modified cells with wild-type counterparts rescues the functional deficit by allowing access to operational proteins in neighboring cells. We propose that this rescue lessens the deleterious effects of mutations in genes related to metabolism. For such processes, a cell should not be considered as the unit under selection in somatic evolution.

## RESULTS

### Screening colorectal cancer cells for connexin expression

Microarray data from seventy-nine CRC lines (23) were analyzed for the expression of connexin (Cx) genes. Expression in a subset of CRC cells was detected for *GJA1* (coding for Cx43), *GJA3* (Cx46), *GJB1* (Cx32), *GJB2* (Cx26), *GJB3* (Cx31), *GJB5* (Cx31.1) and *GJC1* (Cx45) (**Figure 1A)**. Gaussian mixture modeling determined that the distributions of *GJA1, GJA3, GJB1, GJB5* and *GJC1* message among CRC lines were bimodal, so cells could be grouped by high or low expression. In contrast, messages for *GJB2* and *GJB3* were unimodal. Analysis of TCGA datasets confirmed bimodality for *GJA3*, *GJB5* and *GJC1* expression (**Figure S1)**. Single-cell transcriptomics (24) indicated expression of *GJB1*, *GJB2* and *GJB3* in normal colonic epithelial cells (**Figure 1B**). *GJB2* and *GJB3* had previously been identified in normal colorectal epithelium (25, 26). Thus, *GJB2* and *GJB3* are constitutively expressed in CRC cell lines, whereas *GJA1*, *GJA3* or *GJC3* were found in a subset of lines due to mutations or stable methylation differences in promoter regions. Two-by-two table analysis for bimodally-distributed connexin genes indicated positive correlations between *GJA1* and *GJC1*, and negative correlations for pairs *GJB1*-*GJA1*, *GJB1*-*GJB5* and *GJB1-GJC1* (**Figure 1C;** see also principal component analysis **Figure S2**).

**Figure 1:**
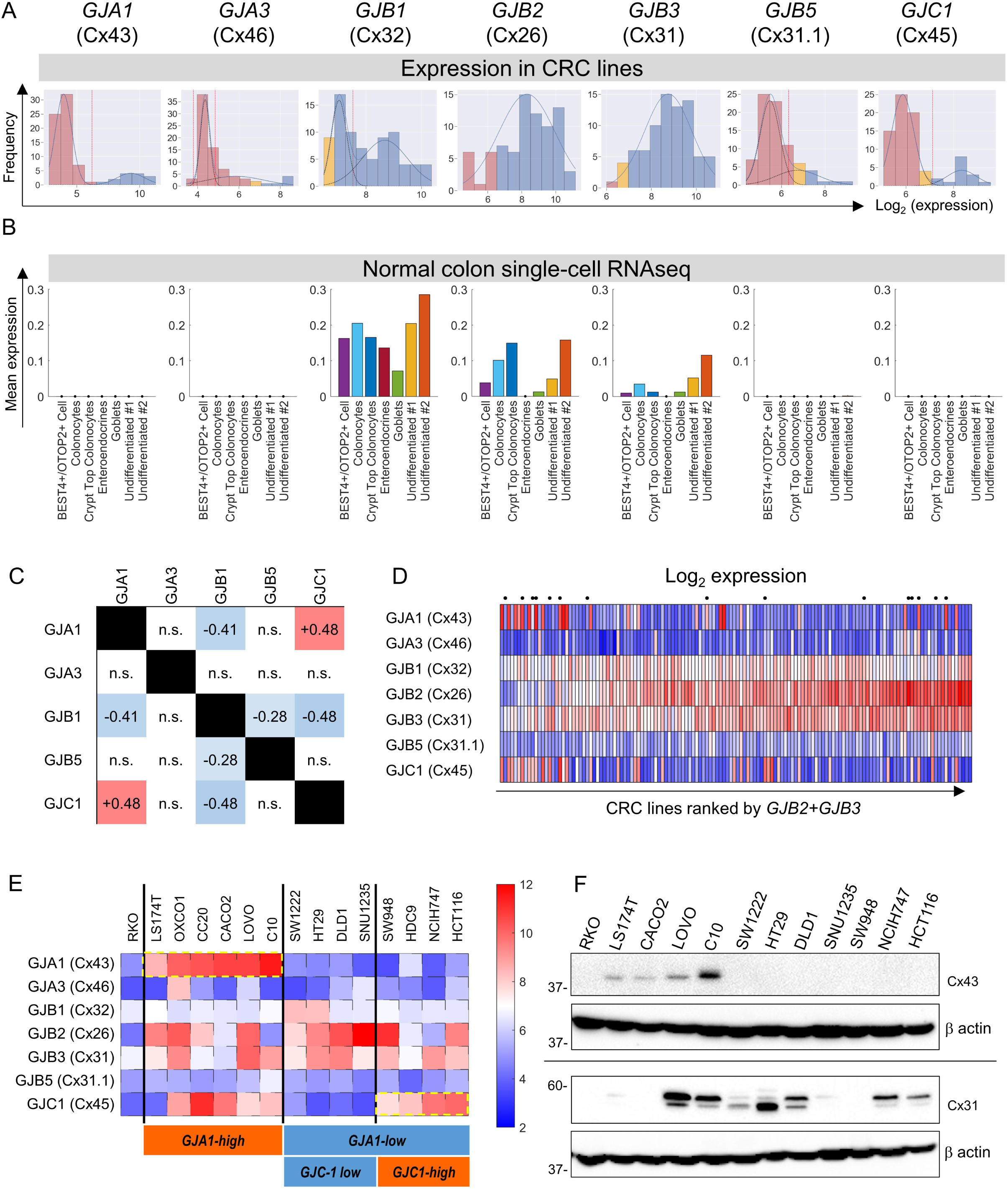
*Connexin isoform expression in CRC cells.* (A.) Microarray data from 79 CRC cell lines analyzed for message level of seven connexin genes. Frequency distributions for log_2_-transformed data. A Gaussian mixture modelling based analysis (GMMchi) is used to determine whether the distributions are bimodal or unimodal. Vertical red line is the (cut-off threshold separating low/high groups. Pink bars refer to background levels and yellow bars refer to near-background levels, based on a separate analysis of the overall pattern of gene expression observed in the cell lines. The cut-off thresholds for the difference between low and high expression are: 2^6.2^, 2^4.9^, 2^7.5^, 2^6.3^, and 2^7.1^ for *GJA1*, *GJA3*, *GJB1*, *GJB5*, and *GJC1*, respectively. (B.) Analysis of single-cell RNAseq datasets for normal colon obtained from the GSE116222 dataset available at the Gene Expression Omnibus (https://www.ncbi.nlm.nih.gov/geo/query/acc.cgi?acc=GSE116222). Bars show mean expression levels by cell type. (C.) Two-by-two table shows correlation between bimodally distributed connexin genes. Numbers refer to correlation coefficient for significant (p<0.05) gene pairs (Fisher’s exact test). (D.) Log_2_-transformed expression data ranked by the sum of *GJB2* and *GJB3* expression. Cells selected for further studies are indicated by a dot above the heatmap. (E.) Heatmap replotted for the selected 15 cell lines, grouped by *GJA1* and *GJC1* expression, relative to threshold determined from GMM analysis. (F.) Western blots for Cx43 and Cx31, showing agreement between protein levels and gene expression profiles.

**Figure 1D** shows the expression of connexin genes, ranked by combined *GJB2* and *GJB3* message. Fourteen lines were selected for further functional measurements (**Figure 1E**). Since *GJA1* expression accounted for considerable variation, cell lines were categorized as *GJA1*-high and *GJA1*-low. The *GJA1*-low group could be subdivided as *GJC1*-low and *GJC1*-high. This classification was confirmed at protein level using antibodies against Cx43 (*GJA1*) and Cx31 (*GJB3*) (**Figure 1F**). RKO cells were chosen as a negative control because of negligible expression of all Cx genes.

### Cx26 (GJB2) channels are a major route for diffusive exchange in CRC cells

Soluble diffusion between cells was measured by FRAP (fluorescence recovery after photobleaching) of calcein (27). A cell in the middle of a coupled cluster was bleached to ∼50% of resting signal; the subsequent recovery of fluorescence due to dye ingress from neighboring cells indicated the degree of coupling. Fluorescence recovery was detected in *GJB2*-expressing SNU1235 but not in RKO cells (**Figure 2A**). FRAP recordings were quantified in terms of a permeability constant (**Figure 2B**), and the role of Cxs was confirmed from the inhibitory effect of carbenoxolone (100 μM), a broad-spectrum Cx blocker (**Figure 2C**). Cell-to-cell permeability correlated best with *GJB2* expression (**Table S1**). *GJB2* knockdown ablated functional coupling, whereas siRNA against *GJB3* did not (**Figure 2D**). In the case of *GJA1*-high CACO2 cells, functional coupling was ablated with siRNA against *GJA1* (**Figure 2E**). Coupling correlated with Cx26 protein at cell-to-cell contacts (**Figure 2F**). Cx43 was detectable at cell-to-cell contacts in C10, LOVO and CACO2 but not in RKO, SW1222 or HT29 cells (**Figure 2G**). Ablation of *GJB2* expression by guide RNA (gRNA) or siRNA (**Figure 2H**) eliminated Cx26 immunofluorescence (**Figure 2I**). *GJB2* knockout (KO) clones of DLD1 cells, generated by CRISPR/Cas9 (**Figure 2J**) had dramatically reduced diffusive exchange (**Figure 2K**). In summary, Cx26 channels provide a principal route for cell-to-cell communication in most CRC lines.

**Figure 2:**
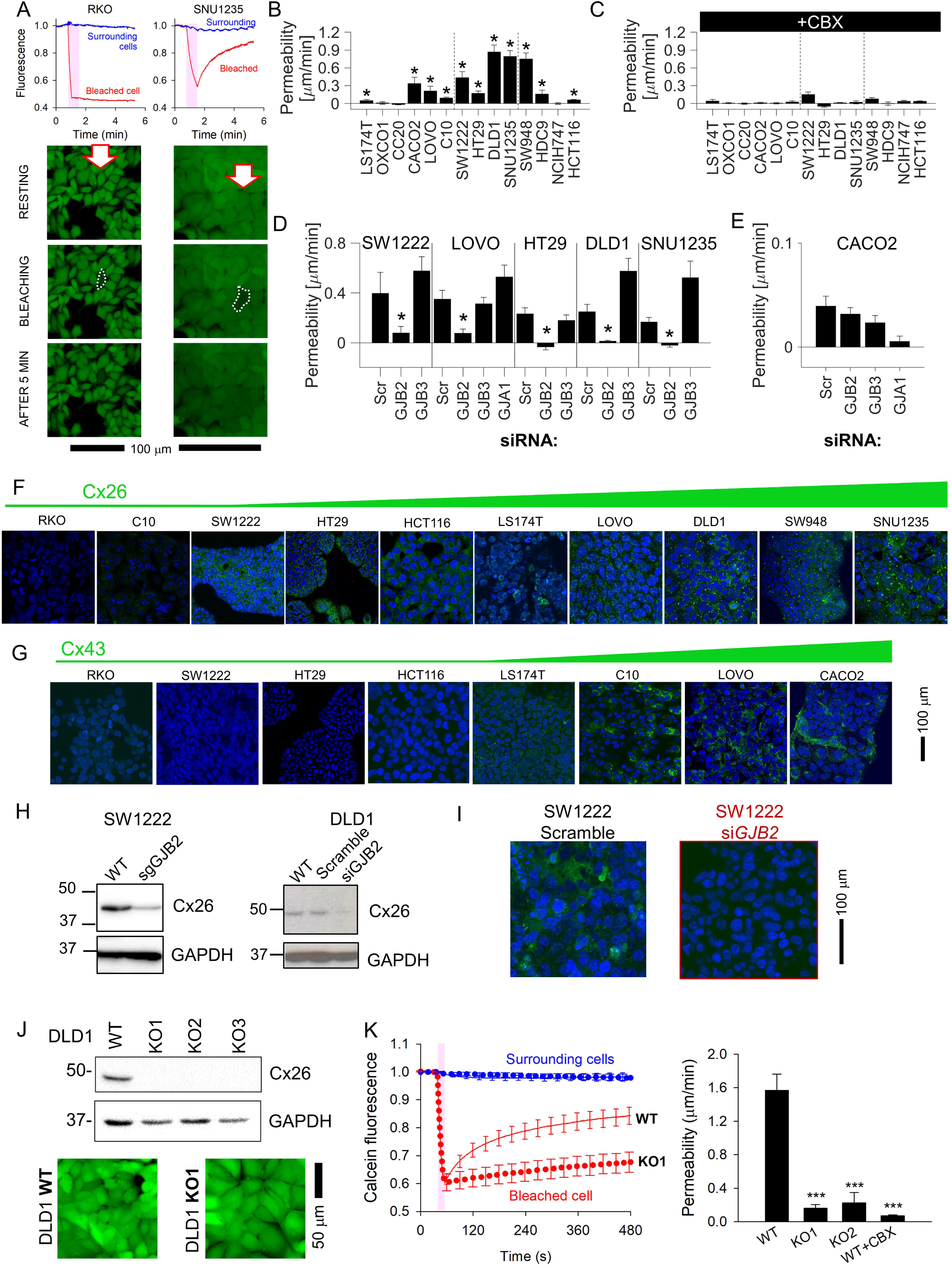
*Connexin isoforms underpinning cell-to-cell coupling in CRC cells.* (A.) FRAP protocol for interrogating cell-to-cell permeability to calcein in RKO cells (connexin-null) and SNU1235 cells (Cx26-positive). Images taken before bleaching (resting), immediately after bleach, and 5 min after bleach. (B.) Permeability (mean±SEM) in CRC monolayers; * denotes significant coupling (t-test). For each experiment, measurements were obtained from at least 5 independently grown monolayers, with multiple technical repeats each. N=15-80 per line. (C.) FRAP measurements repeated in the presence of 100 μM carbenoxolone (CBX). (D/E.) FRAP measurements on cells transfected with siRNA to knockdown *GJA1*, *GJB3* or *GJB2*. * denotes significant decrease in permeability relative to scrambled construct control. (F.) Immunofluorescence in monolayers showing nuclei stained with DAPI (blue) and connexin Cx26 (green), where present. (G.) Immunofluorescence performed with Cx43 antibody (green). Images ranked by increasing connexin signal at cell-to-cell contacts. (H.) Confirmation that sgRNA or siRNA against *GJB2* decreases the expression of Cx26 in DLD1 and SW1222 cells. (I). siRNA knockdown of *GJB2* eliminates Cx26 immunofluorescence signal at cell-to-cell contacts in SW1222 cells. (J.) Blot for Cx26 in DLD1 cells and knockout (KO1-3) clones, and confocal image of Calcein-loaded monolayers established from for WT or KO1 cells. (K.). Confluent DLD1 *GJB2* KO (clone 1) monolayers had substantially reduced cell- cell connectivity, as determined by FRAP. Mean±SEM of 20 cells from 3 monolayers for each genotype.

### Small cytoplasmic molecules equilibrate across coupled CRC monolayers

FRAP-based measurements can identify the Cx channels that underpin permeability but cannot predict the extent to which solutes equilibrate at steady-state. This parameter was evaluated from the degree of fluorescent dye exchange in co-cultures prepared from two populations of cells loaded with spectrally distinct CellTracker fluorophores. The diffusive properties of CellTracker dyes, determined by FRAP (**Figure 3A**), correlated inversely with molecular weight (**Figure 3B**). CellTracker Violet and Orange were selected for co-culture experiments (**Figure 3C**). Monolayers were imaged sequentially for Violet or Orange in confocal mode. In control monolayers prepared with one dye only, fluorescence bleed-through between channels was shown to be minimal (**Figure 3D**). CellTracker Violet and Orange fluorescence images were pseudo-colored as blue and red respectively, such that voxels containing both dyes (i.e. diffusive exchange) appear purple (**Figure 3D**).

**Figure 3:**
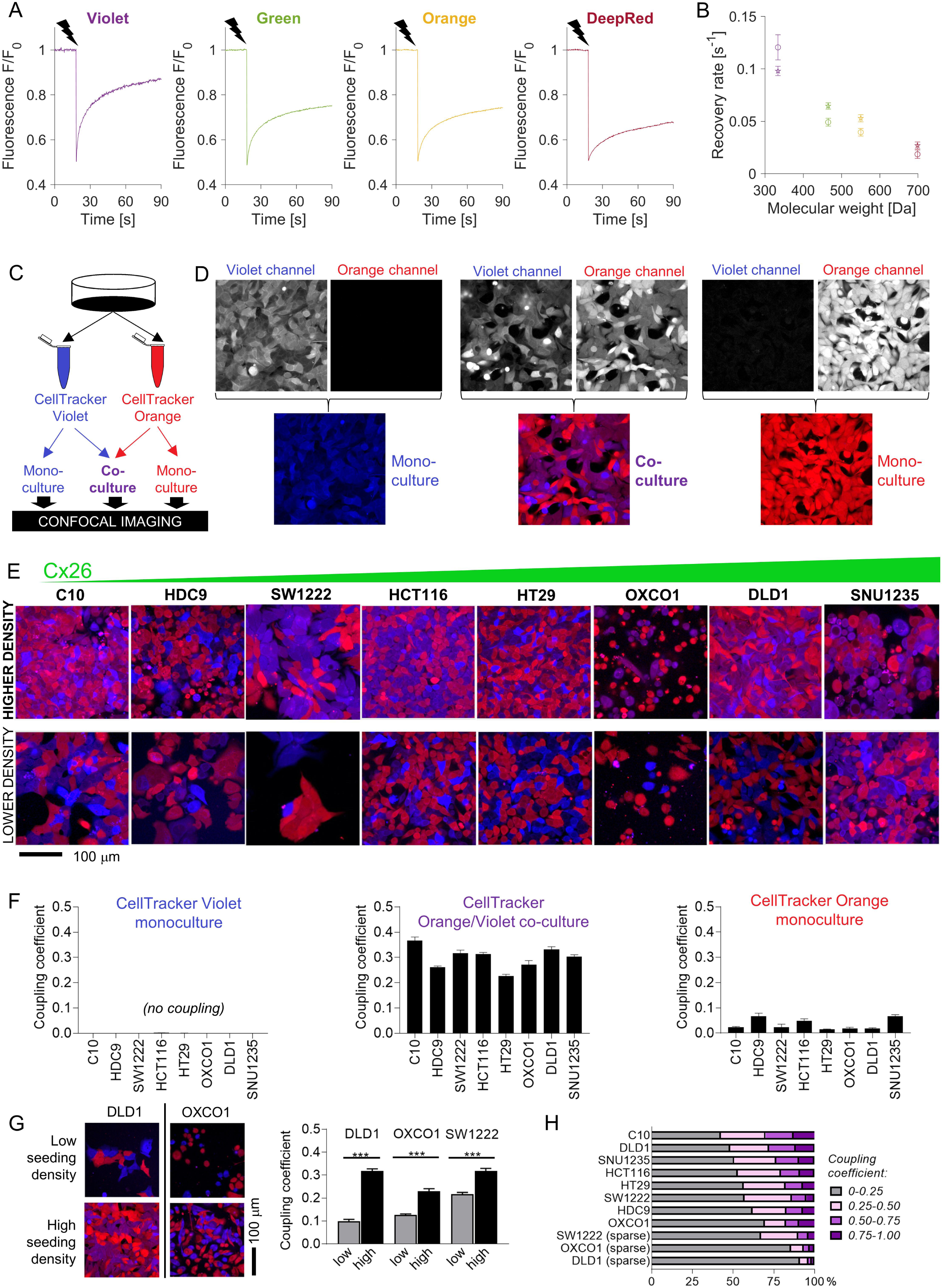
*Fluorescent molecules equilibrate between coupled cells: imaging.* (A.) Representative time courses of FRAP protocol for measuring cytoplasmic diffusivity of CellTracker dyes in DLD1 cells in sparse culture. (B.) Mean recovery rate constant as a function of the molecular weight of the CellTracker dye. Mean±SEM; N=20-25 DLD1 cells (star symbol), 7-15 LOVO cells (circles). (C.) Schematic for preparing co-cultures or monocultures loaded with CellTracker dyes. (D.) Confocal imaging of DLD1 monolayers. CellTracker orange is pseudocolored red and CellTracker Violet is pseudocolored blue; mixing produces purple appearance. (E.) Images of co-cultures from eight CRC lines, ranked by increasing *GJB2* message, acquired from confluent and low-confluency regions. (F.) Quantification of dye exchange in terms of coupling coefficient. Note that mono-cultures produce no (Violet) or very low (Orange) coupling coefficients. Note that low coupling coefficients in some CellTracker Orange monocultures was due to autofluorescence detected on the violet channel in the absence of CellTracker Violet. Mean±SEM of N=20-30 fields of view from 4-5 biological repeats; co-cultures and mono-cultures paired. (G.) Comparison of coupling in cell lines plated at low density to reduce the incidence of cell-to-cell contacts. Pairwise significance testing by t-test: *** is P<0.001. (H.) Frequency distribution of coupling coefficient ranked by decreasing coupling strength.

In eight CRC lines spanning the range of *GJB2* expression, dye exchange was confirmed, particularly in areas of high cell confluency (**Figure 3E**), which points to the role of cell-to-cell contacts. In monocultures, the coupling coefficient was very low, as expected from a negative control. Co-cultures, in contrast, produced significant coupling coefficients (**Figure 3F**). Strikingly, there was no correlation between FRAP- measured permeability and coupling coefficient determined at steady-state. Thus, even monolayers with low-conductance gap junctions can produce a uniform distribution of diffusible substances at steady-state.

Some degree of dye leakage from cells is inevitable during long-term culture, but this is unlikely to result in the transfer of dye into neighboring cells. Firstly, CellTracker dyes become membrane-impermeable inside cytoplasm, thus any molecules that leak across damaged membranes cannot enter neighboring cells. Secondly, the volume of extracellular space is far greater than intracellular volume, thus any released dye would become vanishingly diluted outside cells. To confirm that the dye-mixing was due to exchange across cell-to-cell contacts, co-culture experiments were repeated on cells seeded at very low density to reduce the incidence of cell-to-cell contacts (**Figure 3G**). These produced significantly lower coupling coefficients (**Figure 3H**).

Dye exchange was tested further using flow-cytometry (**Figure 4A**). Co-cultures were prepared using various pairs of spectrally-resolvable CellTracker dyes. The gain of detection channels was calibrated using mono-cultures prepared using one dye only. After gating-out duplets, fluorescence was measured on two detection channels. DLD1 cells grown as mono-cultures emitted fluorescence detected on one channel only (**Figure 4B**). Co-cultures, in contrast, produced a cluster of cells that emitted fluorescence in both channels, indicating dye exchange during culture. This can be visualized by overlaying pseudo-colored density histograms for co-cultures (as green) against the two mono-cultures (as red or blue). To confirm that dye exchange had taken place whilst cells were connected as part of an intact monolayer, control experiments mixed two suspensions of cells harvested from distinctly labelled mono- cultures immediately prior to cytometry. These experiments produced two separate clusters, confirming that dual labelling arises only in intact co-cultures (**Figure 4C**). Similar experiments were performed in other CRC cells (**Figure 4D-G**). The extent of dye exchange between coupled cells was ∼60% (DLD1 or HT29 cells; **Figure 4H**).

**Figure 4:**
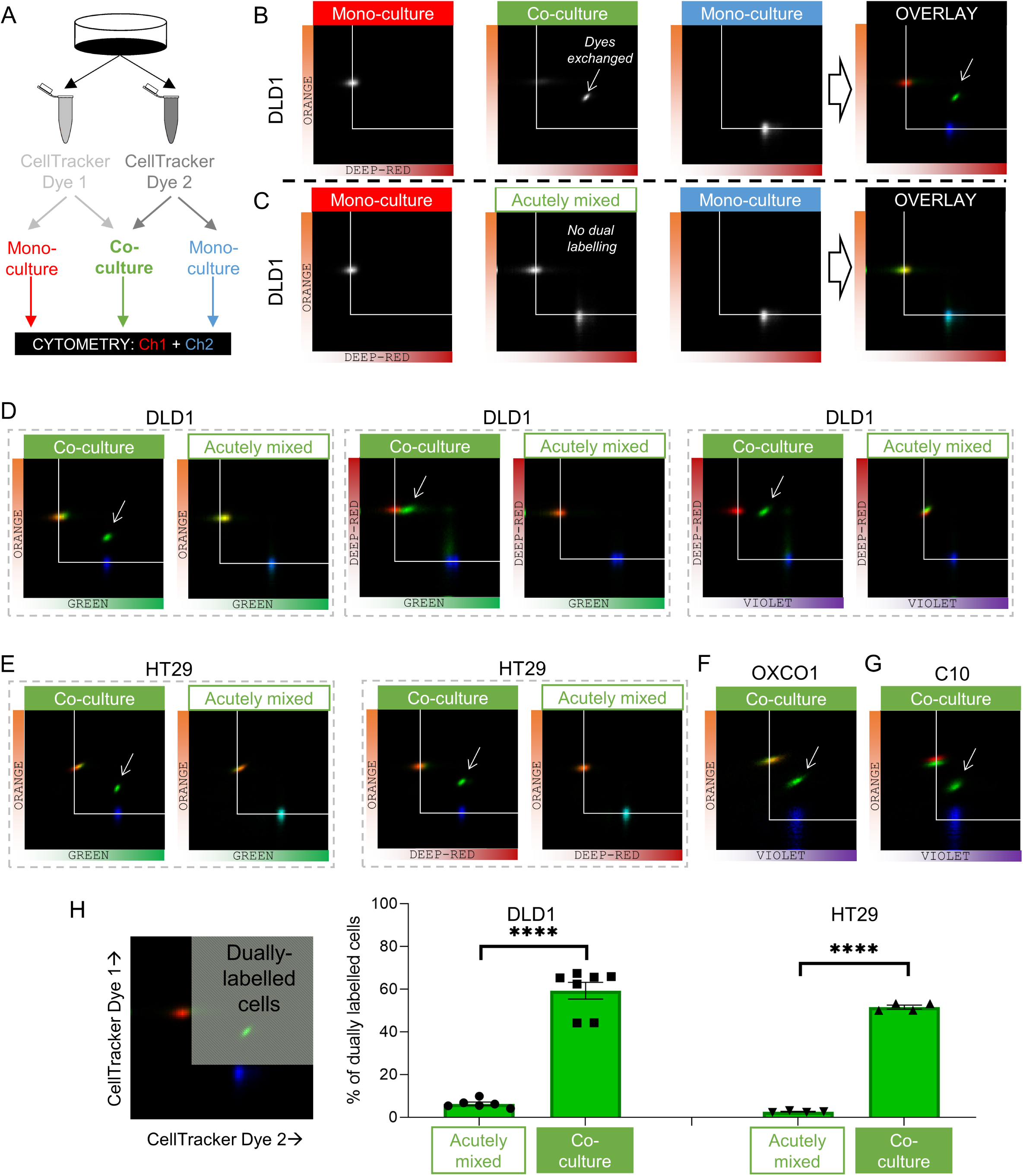
*Fluorescent molecules equilibrate between coupled cells: cytometry.* (A.) Schematic for producing monocultures or co-cultures with pairs of CellTracker dyes. (B.) Cytometry of DLD1 monocultures grown from cells loaded with either DeepRed or Orange, or co-cultures grown from 1:1 mix of cells loaded with DeepRed and Orange. Logarithmic axes showing signal on the relevant detection channels. CellTracker Orange monocultures emit signal on the orange detection channel only; CellTracker DeepRed monocultures emit signal on the DeepRed detection channel only; co-cultures emit fluorescence on both detection channels (indicating that dye exchange had taken place). Overlay shows pseudo-colored bivariate density maps: red for Orange mono-cultures, blue for Deep-Red mono-cultures and green for co-cultures. Arrow points to cluster showing evidence for dye exchange. (C.) Flow Cytometry of DLD1 cells that were grown as monocultures labeled with either DeepRed or Orange. Aliquots of DeepRed- and Orange-labelled cells were mixed prior to flow cytometry to test for acute dye exchange. Analyses show no evidence for dye exchange between acutely mixed cell suspensions. (D.) Cytometry of co-cultured or acutely mixed DLD1, (E.) HT29, (F.) OXCO1 or (G) C10 cells loaded with various pairs of CellTracker dyes. (H.) Quantification of the fraction of cells that had exchanged CellTracker dyes. Region thresholds defined by 95^th^ percentile of signal detected on channels. Significant dye exchange took place only in co-cultured cells but not acutely mixed cells. Significance testing by t-test: **** is P<0.0001.

In summary, coupled CRC cells can freely exchange solutes of <1 kDa across connexin channels. Thus, small metabolites are expected to equilibrate across the continuous cytoplasmic compartment. The next series of experiments sought evidence that diffusive exchange can functionally rescue cells carrying a genetic inactivation of a specific metabolite-handling process. This was tested in CRC lines that are connected primarily by one connexin isoform so that connectivity could be inactivated by targeting a single connexin gene (i.e. *GJA1*-low cells.

### Coupling rescues genetically inactivated trans-membrane Na^+^/H^+^ exchange

The Na^+^/H^+^ exchanger NHE1, coded by *SLC9A1*, is a prominent regulator of intracellular pH (pHi), expressed at high levels in many CRC cells, including HCT116. Genetic inactivation of NHE1 affects pHi control and is detrimental to cellular physiology (28). If gap junctions were able to conduct a sufficient flow of H^+^ ions between coupled cells, then it is conceivable that an *SLC9A1*-deficient cell could have its pHi-regulatory needs serviced by a neighboring wild-type cell. To test this, two *SLC9A1* KO clones were generated using CRISPR/Cas9 (**Figure 5A**). Confirmation that these KO cells lack NHE1 activity was sought in monolayers loaded with the pH- reporter dye cSNARF1. In wild-type (WT) HCT116 cells, NHE1 activity was measured from the rate of pHi recovery following an acid-load (ammonium prepulse). Experiments were performed in the absence of CO_2_/HCO_3_^-^ to eliminate HCO_3_^-^ dependent transporters. NHE1 activity was confirmed from the inhibitory effect of cariporide (**Figure 5B**). KO monocultures, in contrast, produced no cariporide- sensitive pHi recovery. To test if pHi regulation in KO cells could be rescued by coupling onto WT cells, co-cultures were established, wherein KO cells were identified by transfected GFP fluorescence (**Figure 5C**). pHi recovery, following an ammonium prepulse, was tracked simultaneously in WT and KO cells (**Figure 5D**). Strikingly, KO cells showed evidence for pHi recovery when co-cultured with WT neighbors, despite the absence of *SLC9A1* expression. WT cells in co-culture produced slower pHi recoveries than in monocultures because of the additional burden of having to service KO cells. Similar observations were made with both KO clones. These experiments illustrate phenotypic blurring: apparent function is more homogenous than the underlying genotype.

**Figure 5:**
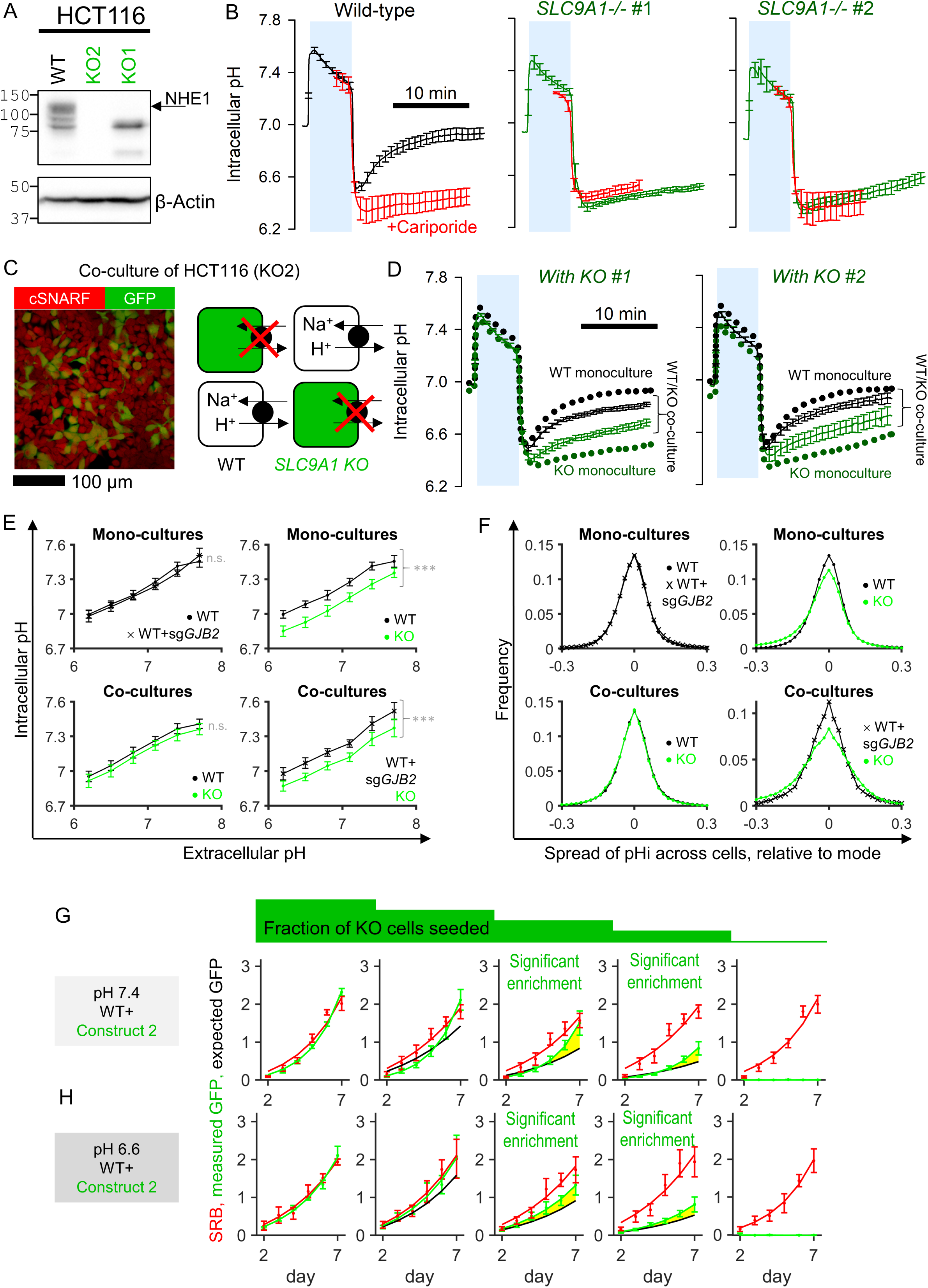
*Diffusive coupling rescues genetically-inactivated Na^+^/H^+^ exchange function.* (A.) Western blot for NHE1, product of *SLC9A1* in wild-type HCT116 and two KO clones. gRNA for KO1 produced a truncated protein, whereas gRNA for KO2 produced complete ablation of expression. (B.) Functional confirmation of genetic inactivation of *SLC9A1*. Measurements of intracellular pH (pHi) in HCT116 monolayers: following acid-loading by ammonium prepulse (absence of CO_2_/HCO_3_^-^ buffer), NHE1 activity in wild-type (WT) cells mediates recovery of pHi (N=13 biological repeats, with 3 technical repeats each), blocked pharmacologically with cariporide (30 µM; N=6 biological repeats, with 3 technical repeats each). KO clones (N=7-9 biological repeats, with 3 technical repeats each) produce no NHE1 activity. Mean±SEM. (C.) Co-culture of WT and KO HCT116 cells. Cells loaded with cSNARF to stain cytoplasm; GFP fluorescence emitted from KO cells. (D.) Ammonium prepulse performed on co-cultures, separating signal from WT (GFP-negative: black) and KO cells (GFP-positive: green). Mean±SEM (N=9-13 biological repeats, with 3 technical repeats each). For comparison, dotted line shows results from WT or KO monocultures. (E.) Relationship between extracellular and intracellular pH measured in HEPES-MES buffered media (absence of CO_2_/HCO_3_^-^ buffer) for WT cells, *GJB2* knockdown cells, or *SLC9A1* KO cells. Results from clone 1 and clone 2 were not significantly different and pooled together. Measurements were performed for monocultures or co-cultures. Mean±SEM (clockwise: N=6 WT and 6 WT *sgGJB2* monocultures; 6 WT and 7 KO monocultures; 8 WT+KO co-cultures; 6 WT sg*GJB2*+KO co-cultures; each with 6 technical repeats). (F.) Analysis of pHi data from E in terms of frequency distribution of pHi, offset to the mode. (G.) Growth curves for HCT116 cells grown as co-cultures of various ratios of WT and KO2 cells, quantified in terms of GFP fluorescence (KO compartment) and SRB absorbance (total biomass) over 7 days of culture, starting from a seeding density of 2,000 cells/well. Mean±SEM (N=5 per construct, with 4 technical repeats each). Significant enrichment indicates that the KO compartment expanded faster than expected from the SRB curve and seeding ratio (2,000:0, 1500:500, 1000:1000, 500:1500 and 0:2000), indicating that KO cells benefited from coupling onto WT neighbors. Statistical testing by two-way ANOVA; P value reported for difference between measured growth of KO cells (GFP) and prediction growth (SRB, scaled by KO:WT seeding ratio). Significance at P<0.05. Media were at pH 7.4. (H.) Experiments were repeated with media at pH 6.6.

To test how diffusive exchange impacts on steady-state pHi, we performed measurements in cells that had equilibrated in HEPES/MES-buffered medium over a range of extracellular pH (pHe). In WT cells, the pHe-pHi relationship was linear and not affected by knockout of *GJB2* (**Figure 5E**). Steady-state pHi was reduced in KO monocultures, as expected from impaired pHi control. However, the pHi difference between WT and KO cells collapsed in co-cultures. This effect was Cx26-dependent because *SLC9A1*-expressing HCT116 cells that had genetically inactivated *GJB2* were unable to raise the pHi of KO cells in co-cultures (**Figure 5E**). Cx26-dependent diffusive connectivity also reduced cell-to-cell variation in steady-state pHi (**Figure 5F**).

To test if diffusive exchange benefits the growth of *SLC9A1*-deficient cells, co-cultures were established from different seeding ratios of WT and KO cells. Parallel plates were set-up to record growth at various time points. Total biomass (i.e. growth of WT and KO cells) was measured by the sulforhodamine B (SRB) absorbance assay, whereas growth of the KO compartment was determined from GFP fluorescence. GFP and SRB signals were calibrated using time courses obtained from KO mono-cultures, for which the two indices are equivalent. If WT and KO cells had identical survival prospects, then the time courses of GFP and SRB should be stoichiometrically related to the seeding ratio. Experiments were performed in media at pH 7.4 or 6.6 on the basis that the latter should amplify the importance of NHE1 (**Figure 5G/H**). For co-cultures seeded from 25% KO or 50% KO cells, growth of the KO compartment (GFP) exceeded the prediction based on equal survival (**Figure 5G/H**), in both clones (**Figure S3**). Thus, co-cultured *SLC9A1*-deficient cells grow faster than expected, at the expense of coupled WT cells. Since pHi regulation consumes a substantial amount of energy, this effect could be explained by the exploitation of WT resources by KO cells.

In summary, we show that cells lacking an important ion exchanger at the membrane can hijack the equivalent activity from neighboring wild-type cells, provided that the ions are able to diffuse across connexin channels.

### Coupling rescues genetically inactivated glycolytic metabolism

The next test related to glycolysis, a critical metabolic pathway that handles small molecules in the cytoplasm. DLD1 cells were selected for these experiments based on their high glycolytic rate. A recent whole-genome CRISPR-Cas9 screen identified *ALDOA*, a gene coding for the glycolytic enzyme aldolase A, as essential for CRC cell survival under physiological pH (29). Ablation of *ALDOA* using virally transduced gRNAs reduced expression (**Figure 6A**) and glycolytic rate, measured in terms of medium acidification (**Figure 6B**) and lactate production (**Figure 6C**). It was not possible to produce a stable *ALDOA* knockout clone, ostensibly because of the enzyme’s critical role for survival, thus experiments were performed within 6 days after transduction with one of the two gRNA constructs.

**Figure 6:**
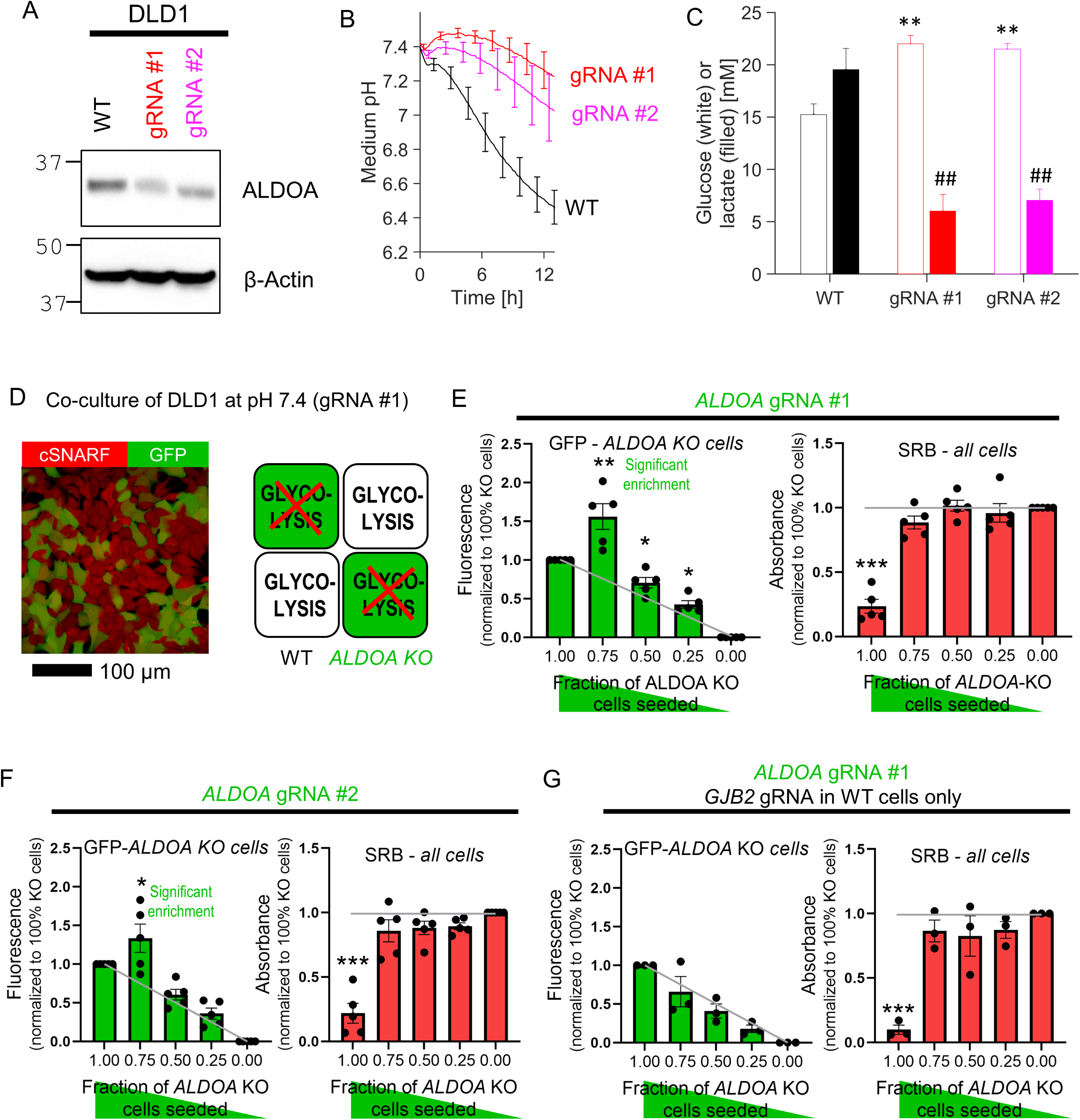
*Diffusive coupling rescues genetically-inactivated glycolysis.* (A.) Western blot for aldolase A (ALDOA) in wild-type (WT) DLD1 cells and cells infected with one of two guide RNA constructs to genetically inactivate *ALDOA*. (B.) Medium acidification measured in confluent DLD1 cells in low-buffer power media (2mM HEPES+MES). WT cells are highly glycolytic, as measured from the acidification time course. Glycolytic rate is greatly reduced by *ALDOA* ablation. Mean±SEM (N=4 plates, each with 3 technical repeats). (C.) DLD1 cells cultured for 4 days in CO_2_/HCO_3_^-^ buffered media. Media collected at end-point were assayed for glucose and lactate. Mean±SEM (N=3 plates, each 3 technical repeats pooled per assay). (D.) Co-culture of WT and *ALDOA*-deficient DLD1 cells. Cells loaded with cSNARF-1 to indicate cytoplasm; GFP fluorescence emitted from gRNA-infected cells. (E). GFP fluorescence and SRB absorbance after 6 days of culture of various seeding ratios of WT and *ALDOA*-deficient cells infected with construct #1: 2000:0, 1500:500, 1000:1000, 500:1500 and 0:2000. Left: grey line shows expected GFP signals if growth of WT and *ALDOA*-deficient cells were no different. Right: gray line shows expected SRB signal if growth of WT and *ALDOA*-deficient cells were no different. Significance testing by t-test relative to gray line. Mean±SEM (N=5 plates, 6 technical repeats each). (F.) Experiment repeated with construct 2. Mean±SEM (N=5 plates, 6 technical repeats each) (G.) Experiment repeated after *GJB2* gRNA treatment to knockout *GJB2* in WT cells, i.e. inactivating connexin coupling between WT and *NDUFS1*-deficient cells. Mean±SEM (N=3 plates, 6 technical repeats each).

*ALDOA*-deficient DLD1 cells grew four-times slower than wild-type cells (**Figure S4**), likely due to compromised provision of glycolytic ATP and build-up of fructose-1,6-bisphosphate, the substrate for aldolase A (**Figure S4**). These metabolites are expected to diffuse freely between coupled cells, thus any fructose- 1,6-bisphosphate that accumulates in an *ALDOA*-deficient cell should diffuse to neighboring cells for processing, whereas ATP will diffuse in the opposite direction. Diffusive rescue would allow *ALDOA*-deficient cells in co-culture with wild-type cells to grow faster than expected. To test this, co-culture experiments were performed for various ratios of WT and GFP-labelled *ALDOA*-deficient cells (**Figure 6D**). After 6 days of culture at pH 7.4, GFP fluorescence (a readout of *ALDOA*-deficient cells) and SRB absorbance (total cell biomass) were recorded sequentially. If *ALDOA*-deficient cells received no benefit from being coupled onto WT cells, then the end-point GFP signal would be proportional to the initial seeding density (**Figure 6E** grey line). However, GFP-labelled cells became relatively enriched in co-cultures with WT cells, which was most prominent at 3:1 seeding ratio. Thus, the survival of *ALDOA*-deficient cells improved when they had diffusive access to WT cells. A similar effect was seen using the other gRNA (**Figure 6F**). Rescue of *ALDOA*-deficient cells in co-culture with WT cells was absent when *GJB2* was knocked-out in WT cells with gRNA (**Figure 6G**), which indicates a critical role for Cx26-mediated metabolite exchange in this rescue.

In summary, we demonstrate that the genetic ablation of a metabolite-handling cytoplasmic enzyme in one cell can be rescued by diffusive access to the required catalysis in neighboring cells via connexin channels. Thus, despite major differences in genotype, the apparent phenotype of KO and WT cells is more similar.

### Coupling rescues genetically inactivated mitochondrial respiration

The third test for metabolite exchange used cells with genetically inactivated mitochondrial respiration on the basis that this organellar process influences the levels of diffusible substances in cytoplasm, including ATP. SW1222 cells were selected on the basis of their high respiratory rate (**Figure 7B**), which could be genetically inactivated by knockout of *NDUFS1*, a gene coding for a component of complex I (**Figure 7A**) (29). Oxidative phosphorylation was blocked in *NDUFS1* KO cells, and a compensatory increase in glycolytic rate was observed (**Figure 7B**). *NDUFS1*- deficient cells grew three times slower than WT cells (**Figure S5**), likely due to ablated mitochondrial metabolism, critical for delivering substrates and energy. To test if junctional coupling could rescue *NDUFS1*-deficient cells, WT and GFP-labelled KO cells were co-cultured in medium at pH 7.7, alkaline conditions that facilitate a full glycolytic rate (**Figure 7C**). If KO cells received no benefit from being coupled onto WT cells, the end-point GFP signal would be proportional to the seeding ratio. However, co-culture with WT cells promoted the growth of GFP-labelled KO cells beyond the expected level, particularly for seeding ratios of 3 KO:1 WT, indicating a functional rescue (**Figure 7D**). This effect related to the density of cell-to-cell contacts, because KO cells could no longer receive a growth benefit in sparse co-culture with WT cells (**Figure 7E**). Furthermore, the rescue was linked to coupling involving Cx26, as co-culture with WT cells that had genetically knocked-down *GJB2* by siRNA yielded no benefit to co-cultured *NDUFS1*-deficient cells (**Figure 7F**). Ultimately, the rescue effect of WT cells relates to oxidative phosphorylation, which requires oxygen. Thus, inactivating this process under hypoxic conditions is predicted to attenuate the benefit to co-cultured *NDUFS1*-deficient cells. Indeed, this was the case in co-cultures incubated in 2% O_2_ (**Figure 7G**).

**Figure 7:**
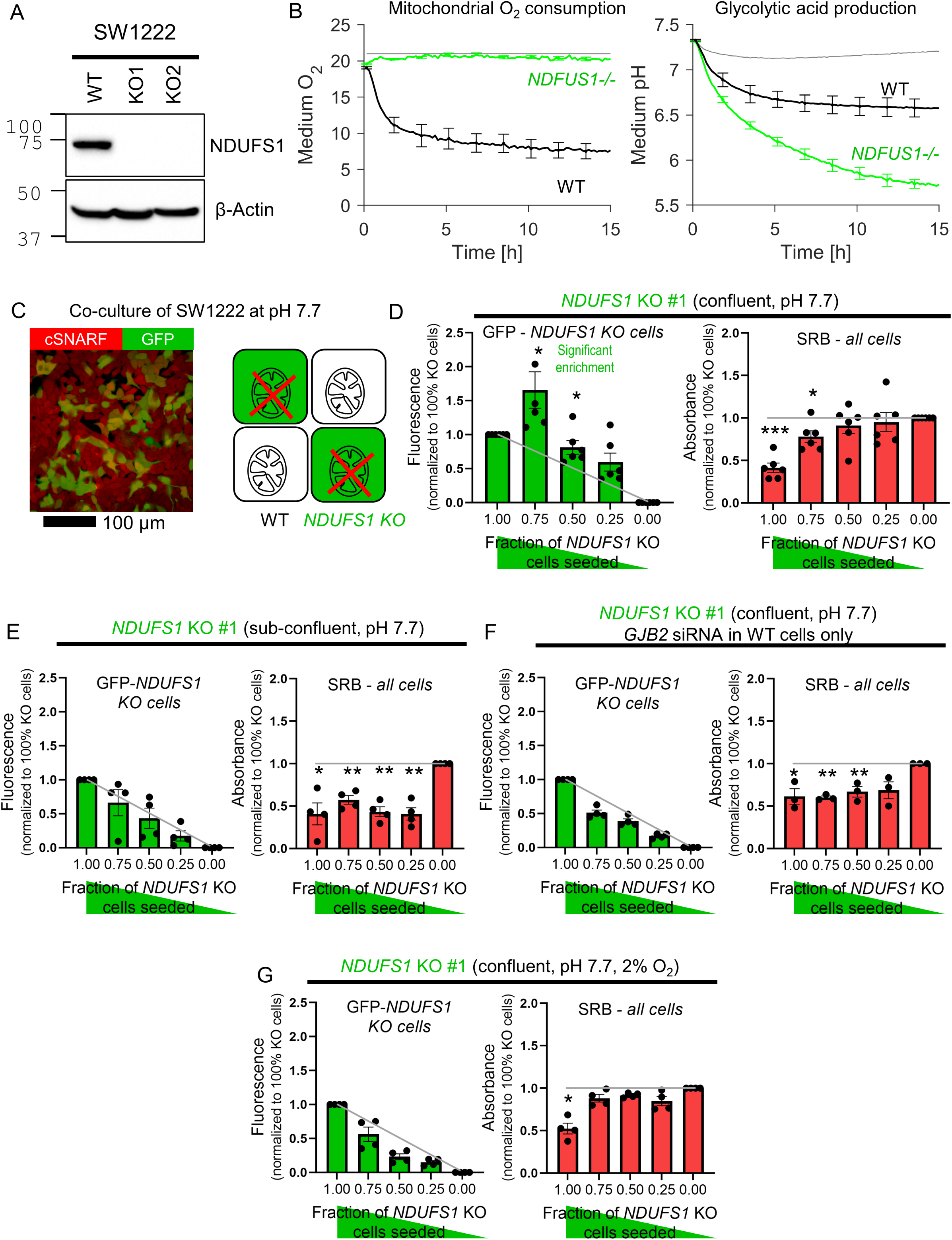
*Diffusive coupling rescues genetically-inactivated mitochondrial respiration.* (A.) Western blot for NDUFS1 in wild-type (WT) SW1222 cells and *NDUFS1* KO clones established using one of two guide RNAs. (B.) Fluorimetric measurements of oxygen consumption and acid production (assays of respiratory and glycolytic rates) in WT and *NDUFS1* KO SW1222 cells. Mean±SEM (N=3 plates, 3 technical repeats each). (C.) Co-culture of WT and *NDUFS1* KO cells. Cells loaded with cSNARF-1 to indicate cytoplasm; GFP fluorescence emitted from KO cells. (D) GFP fluorescence and SRB absorbance after 6 days of culture of various seeding ratios of WT and *NDUFS1*- deficient cells infected with construct #1: 2000:0, 1500:500, 1000:1000, 500:1500 and 0:2000. Left: grey line shows expected GFP signals if growth of WT and *NDUFS1*- deficient cells were no different. Right: gray line shows expected SRB signal if growth of WT and *NDUFS1*-deficient cells were no different. Media pH set to 7.7. Significance testing by t-test relative to gray line. Mean±SEM (N=4 plates, 4 technical repeats each). (E.) Experiment repeated at lower (1000/well) seeding density to reduce incidence of cell coupling. Mean±SEM (N=4 plates, 6 technical repeats each). (F.) Experiment repeated with WT cells *GJB2* KD via siRNA, i.e. inactivating connexin coupling between WT and *NDUFS1*-deficient cells. Mean±SEM (N=3 plates, 6 technical repeats each). (G.) Experiment repeated under hypoxic conditions, which suppresses mitochondrial function. Mean±SEM (N=4 plates, 6 technical repeats).

In summary, we show that genetically inactivated mitochondrial metabolism in one cell can be rescued by coupling onto a wild-type cell. This demonstrates that apparent differences in phenotype can be reduced by diffusive coupling, even if the inactivated process occurs in a subcellular compartment.

## DISCUSSION

This study demonstrated that CRC cells establish a multi-cellular cytoplasmic continuum through which small molecules can diffuse and equilibrate. The diffusive coupling reduces gradients of solutes, blurring differences in metabolite levels between neighboring cells. Solute exchange was demonstrated using fluorescent dyes in terms of their permeability and degree of mixing at steady-state. Permeability, as measured by FRAP, showed variation among CRC cell lines, which correlated best with *GJB2* expression and Cx26 immunoreactivity at cell-to-cell contacts. The role of this connexin isoform in producing functional conduits was inferred from the effect of genetic knockdown on permeability. *GJB2* is expressed in normal colorectal epithelium (30), and the unimodal distribution of its message level among CRC lines suggests that transformed cells mostly retain its expression. In addition to *GJB2*, some CRC cells express other isoforms such as *GJA1* and *GJC1*. Mutations or stable epigenetic changes that activate the expression of *GJA1* or *GJC1*, establishing additional conductance pathways between cells. Intriguingly, the exact magnitude of permeability did not predict the degree to which solutes can be equilibrated between cells at steady-state. This is because even weakly coupled confluent monolayers were able to exchange dyes, if allowed sufficient time. Thus, the phenomenon of solute equilibration across coupled cancer cells is likely to be significant even in cells with low connexin expression.

Various enzymes and transporters handle diffusible molecules which may cross connexin channels and equilibrate between cells. If diffusion takes place, the functional consequences of a genetic inactivation of a transporter or enzyme in one cell could be rescued by access to functional proteins in neighboring cells. This prediction was tested using three examples of biologically important processes that handle small molecules. The three chosen processes were localized to different sub- cellular domains: at the plasma membrane (NHE1, coded by *SLC9A1*, handling intracellular H^+^ ions), in the cytoplasm (aldolase A, coded by *ALDOA*, handling the glycolytic intermediate fructose-1,6-bisphosphate) and in mitochondria (NADH:ubiquinone oxidoreductase core subunit S1; *NDUFS1*, part of complex I). In all three cases, genetic inactivation resulted in a functional defect measured in monocultures, impaired pHi regulation (*SLC9A1*-deficient cells), blocked glycolysis (*ALDOA*-deficient cells) or inactivated mitochondrial respiration (*NDUFS1*-deficient cells). The CRC cells selected for these experiments, HCT116, DLD1 and SW1222, manifested a reliance on NHE1 for pH control, a highly glycolytic phenotype, and a strong dependence on OXPHOS, respectively. Importantly, these cells are *GJA1*- negative, which ensures that Cx43 does not contribute a secondary conductance pathway in parallel to Cx26. This made it easier to control conductance experimentally by targeting one type of connexin (*GJB2*).

For all three examples of metabolic processes, Cx26-dependent connectivity rescued the functional deficit arising from genetic inactivation, provided that the affected cells can access normal proteins in neighboring wild-type cells. These co-culture experiments model the consequences of a spontaneous mutation arising in a cell within a coupled network. Thus, *SLC9A1*-deficient cells can still manifest normal pHi control, if this is serviced by the wild-type protein in neighboring cells and H^+^ ions can diffuse across connexin channels. The survival of *ALDOA*-deficient cells is improved if these can access wild-type cells with normal glycolytic pathways and the relevant metabolites diffuse across connexin channels. Similarly, *NDUFS1*-deficient cells are rescued if these can access normal mitochondrial function in neighboring cells. Diffusive rescue was attributed to Cx26 channels but, in principle, any connexin channel could serve this role. These observations are significant for somatic evolution because cells bearing mutations in metabolite-handling genes may nonetheless fare normally and evade negative selection, provided they are able to connect to a wild-type neighbor. We speculate that this is an important reason why genes described as ‘essential’ by *in vitro* knockout screens may still incur loss-of-function mutations in human cancers without reducing their growth. Such genes would manifest their essentiality when mutated in confluent monocultures *in vitro*, but not necessarily when mutated spontaneously in human tumors, wherein the coupled cellular network compensates for the local defects in function. In contrast, genes responsible for processes that do not handle diffusible solutes cannot benefit from connexin-mediated coupling; for example, genes coding for ribosomal subunits that are too large to cross connexin channels or genes coding for neoantigens that are, by design, confined to the mutation-carrying cell. Inactivating mutations in such genes may be exceptionally rare in cancers because there are no tangible ways for compensating the functional deficit (10).

Our findings emphasize the importance of measuring phenotypic variation, particularly in instances relating to diffusible molecules, as this may not overlay with the more compartmentalized genotypic landscape. In cases where phenotypic differences are reduced as a result of connexin connectivity, it is not appropriate to consider a single cell as the unit under selection. Intriguingly, connexin genes are very rarely mutated in cancer (10). We speculate that this essentiality reflects a critical role of cell-to-cell coupling for cancer growth, and an opportunity for therapeutic interventions.

## MATERIALS AND METHODS

### Cell lines and culture

Human colorectal cancer cell lines were provided by Professor Walter Bodmer. Cells were cultured using DMEM (Sigma-Aldrich, D7777) supplemented with 10% FBS and 1% PenStrep (10 000 U/mL) at 37°C and 5% CO_2_. To adjust medium pH, the content of NaHCO_3_ was varied (31).

### siRNA transfection

Cells were seeded at a density of 200,000 cells/well in a 6 well plate and transfected with either siRNA (20nM), e.g. for *GJB2*, *GJB3* and *GJA1*, or with a scramble non-targeted siRNA (Dharmacon, siGENOME smart pool), using Lipofectamine RNAiMAX (Invitrogen, Cat. No. 2373383). After 72 hours, cells were harvested and seeded for experiments.

### Viral transduction with CRISPR-CAS9 constructs

Gene knockouts were made for DLD1, SW1222 and HCT116 cell lines using DMEM, 10% FBS, 1% Pen/Strep. gRNA sequences were cloned into LentiCRISPR v.2 backbone as previously described (http://genome-engineering.org/gecko/wp-content/uploads/2013/12/lentiCRISPRv2-and-lentiGuide-oligo-cloning-protocol.pdf). Two gRNA sequences were cloned for each gene, using sequences below (Table 1). Concentrated virus aliquots were prepared by the virus production facility at WIMM, University of Oxford. Cells were plated in clear, flat-bottom 6-well plate at a density of 200,000 cells/well and transduced using a 500 μL aliquot of lentivirus carrying the LentiCRISPR v2 construct encoding for a gRNA sequence targeting one individual gene. Polybrene was added at a concentration of 4 µg/mL. The 6-well plate was incubated for 2 days before puromycin (5 μg/mL) was added for selection, and it was incubated for 3 days before the transduced cells were used for further experiments. Single cell clones with stable deletion of *NDUFS1* and *SLC9A1* were obtained in SW1222 and HCT116 cells, respectively. Note that for *ALDOA* gRNA-treated DLD1 cells no stable knock-out clones could be obtained. Therefore, lentivirus pools of knockout cells were mixed populations of cells with different genomic edits and unedited cells. All growth curves were performed within one week of initial viral infection to avoid hypomorph or unedited cells out-competing those that have loss-of-function mutations.

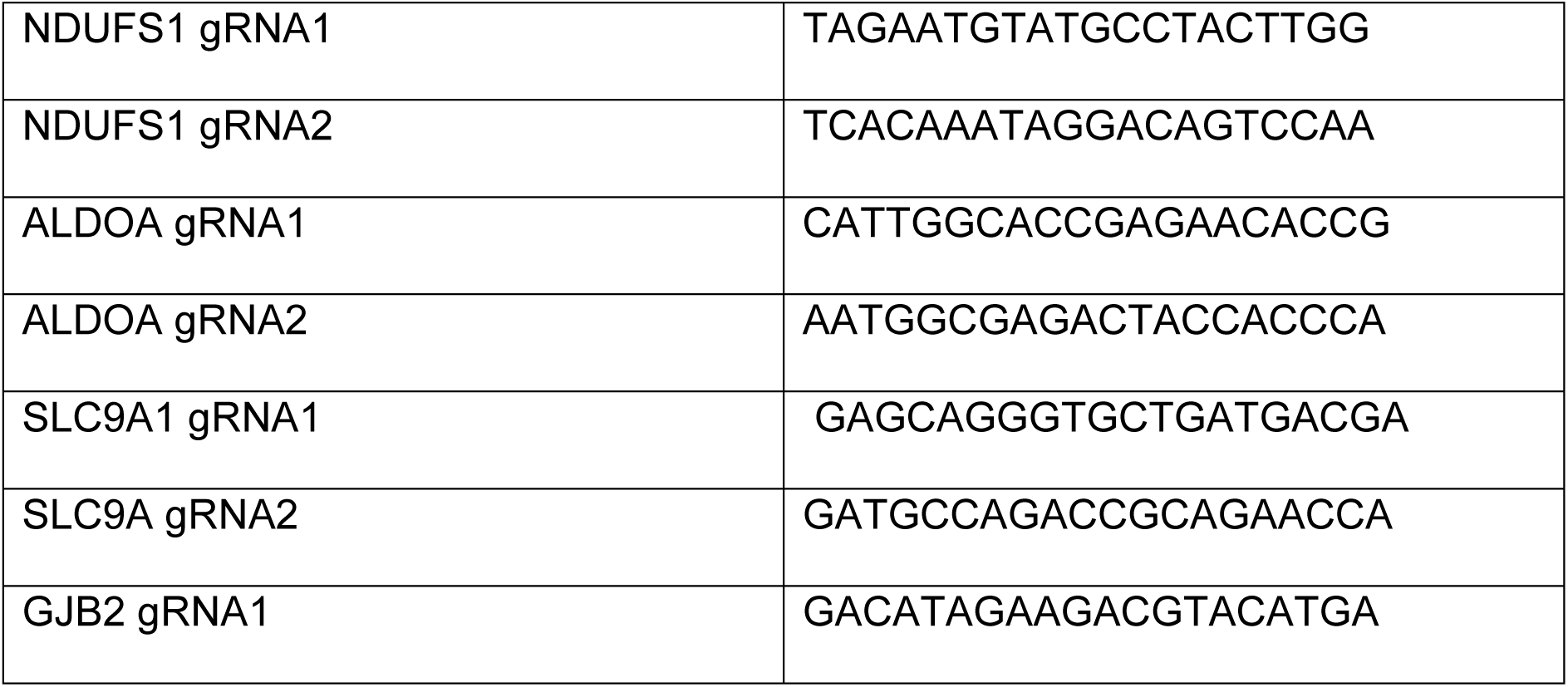

### Immunoblotting

Lysates were prepared with radioimmunoprecipitation assay (RIPA) buffer. Protein concentration in the samples was measured using bicinchoninic acid (BCA) protein assay kit and adjusted using water. Samples were loaded onto a 10% acrylamide gel. The gel was run at 120 V for 90 minutes. Afterwards, membrane transfer was performed at 250mA for 70 minutes. Membranes were incubated with primary antibodies against NDUFS1 (ThermoFisher Cat. No. PA5-22309), Aldolase A (Novus Biologicals, Cat. No. NBP1-87488), NHE1 (BD Biosciences, cat. No.611775), β-actin (Proteintech, Cat. No. HRP-60008) and GAPDH (Proteintech, Cat. No. HRP- 60004) used as a loading control; Connexin26 (Thermo Fisher, Cat no. 13-8100), Connexin 31 (Proteintech, cat. No. 12880-1-AP), Connexin 43 (Thermo Fisher, Cat. No. Cat #13-8300), HRP-conjugated goat anti-rabbit and anti-mouse secondary antibodies were applied. The membrane was visualized using ECL.

### Immunofluorescence

Cells were grown to 50–80% confluency, fixed with 4% paraformaldehyde in PBS (Pierce; Life Technologies) and permeabilized with 0.2% Triton X-100 in PBS. After blocking with 3% BSA in PBS for 1 h, cells were incubated with rabbit antibodies against Cx26 (Proteintech 14842-1-AP) and Cx43 (Cell signalling technology n.3512) for 1.5 hs at room temperature. Cells were then washed and incubated with Alexa Fluor 488 secondary antibody (Life Technologies) for 1 h. Cells nuclei were co-stained with Hoechst 33342 (Life Technologies) applied for 1 min following washing with 1X PBS. The mounting medium used was ProLong Gold Antifade reagent (Invitrogen, Cat. No. P36930).

### Fluorescence Recovery After Photobleaching (FRAP) for measuring coupling

Monolayers in 4-well Ibidi imaging slides were loaded with calcein AM (Invitrogen, C1430) for 10 minutes in Hepes RPMI, replaced to remove unloaded dye. Confluent monolayers were imaged using Zeiss LSM 700 confocal microscope. A cell in the middle of a confluent cluster was selected for bleaching (high-power 488 nm) until fluorescence (488 nm excitation, emission >510 nm) decreased by 50%. Signal was measured in the central cell, neighbors and more remote cells, and normalized to the initial intensity. Recovery was fitted to a mono-exponential to calculate calcein permeability (units: μm/min) from product of the geometric perimeter and area of the central cell, divided by the time constant of fluorescence recovery.

### FRAP for measuring diffusivity

Cells were loaded with Violet (BMQC 5µM, excitation 405 nm, Invitrogen Cat.No. C10094), Green (CMFDA 20µM, excitation 492 nm, Cat.No. C2925), Orange (CMRA 20µM, 555 nm, Cat.No. C34551), or DeepRed (20µM, 630 nm, Cat.No. 34565) for 15 minutes in RPMI Hepes medium. FRAP was limited to a 3x3 μm region of cytoplasm.

### Measuring the exchange of CellTracker dyes by imaging

A suspension of cells was split equally, and each loaded with either CellTracker Violet or Orange for 15 min. Cells were pelleted, washed (PBS), combined in 1:1 ratio, and then seeded onto 4- well imaging slides at 300,000 cells/well. As control, cells were loaded with one type of dye. After 48 h, monolayers were imaged confocally using sequential acquisition that minimizes bleed-through between channels: 405 nm excitation and emission 490- 555 nm for the Violet channel and 555 nm excitation and emission 560-600 nm for the Orange channel. Orange and violet fluorescence were normalized to the mean signal in paired control. The degree of dye mixing was quantified in terms of a coupling coefficient (CC):

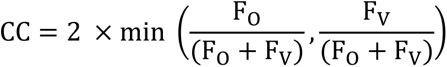

Here, F_O_ and F_V_ are the normalized Orange and Violet fluorescence signals.

### Measuring the exchange of CellTracker dyes by flow cytometry

Cells were loaded with combinations of CellTracker dyes, co-cultured for 48 h and processed for flow cytometry (Attune NXT Analyser). Gain of detection channels was optimized by analyzing monocultures loaded with one dye. Data were presented as pseudo-colored bivariate density plots representing mono-cultures with the low-wavelength dye as blue, mono-cultures with the high-wavelength dye as red, and co-cultures as green.

### Measuring NHE1 activity

Cells seeded on Ibidi 4-well slides were loaded with 10 μM cSNARF1 (Thermo Fisher, Cat. No. C1271, excitation, 555 nm; emission, 580 and 640 nm) for 7 minutes. Cells were then washed by superfusion with HEPES-buffered normal Tyrode containing (in mM): NaCl (135), KCl (4.5), CaCl_2_ (2), MgCl_2_ (1), HEPES (20), glucose (11), pH adjusted to 7.4 with 4 M NaOH heated to 37°C. After 2 min to allow equilibration, the superfusion was switched to ammonium containing Tyrode (in mM): NaCl (105), NH_4_Cl (30), KCl (4.5), CaCl_2_ (2), MgCl_2_ (1), HEPES (20), glucose (11), pH adjusted to 7.4 with 4 M NaOH at 37°C. After 6 min, the superfusate was returned to normal Tyrode. In some experiments, the solutions contained 30 μM cariporide to block NHE1 (Tocris, Cat. No. 5358), as control. In separate experiments, cSNARF1 fluorescence ratio was calibrated using the nigericin method and this curve was used to convert measured cSNARF1 ratio to pHi.

### Measuring resting pHi

Cells were seeded at a density of 80,000 cells/well in Ibidi flat bottom 96 well plates, to produce a monolayer after 24 h. On the measurement day, cells were loaded dually with cSNARF1 (5 μg/ml) and Hoechst 33342 (1:1000) for 15 min, followed by replacement with bicarbonate-free, Phenol Red-free medium based on D5030, containing 10mM HEPES and MES and titrated at pH over the range 6.2 to 7.7. Plates were imaged at Cytation 5 plate reader and each cell, identified from the location of its nucleus, was analyzed for pHi (31). To measure variation of pHi, the frequency distribution was offset to the mode pHi for each experimental condition.

### Measuring medium acidification

Cells were seeded at densities of 2,000 cells per well on clear, flat-bottom 96-well plate (Ibidi). Cells were cultured in bicarbonate-buffered media (D7777, Sigma) set to a pH of 7.4 by adjusting NaHCO_3_ accordingly. The cells were incubated for 4 days at 37°C with 5% CO_2_ before media were removed for glucose and lactate measurements.

### Pentra Assay for glucose and lactate

Cells were seeded onto 96 well plates and grown in DMEM (D7777, Sigma) at pH 7.4. After 5 days, medium was collected, spun to remove residue, and analyzed in a Pentra C400 for glucose and lactate concentrations. Calibrations used standard solutions.

### Measuring metabolic fluxes

Two protocols were performed to measure metabolic rate. The first interrogated glycolytic flux, and the second interrogated glycolysis and respiration simultaneously. For the first, cells were cultured at high density (70,000 cells/well) in flat-bottom, black 96-well plates. To report extracellular pH, media contained 50 µM cSNARF1-dextram. Media, based on DMEM D5030, contained 25mM glucose, 10% FBS, 1% PS, 1 mM pyruvate, 1% glutamax and 2 mM HEPES and 2 mM MES to provide a low but constant buffering power over the pH range studied. Fluorescence was monitored for 17 h using a Cytation 5 device (BioTek, Agilent, Winooski, VT, USA). Excitation was provided by a monochromator, and fluorescence emission was detected sequentially at wavelengths optimized as described previously (32). For the second method, cells were cultured at high density (70,000 cells/well) in flat-bottom, black 96-well plates. To report extracellular pH and O_2_, media contained 2 µM HPTS (8-Hydroxypyrene-1,3,6-trisulfonic acid trisodium salt) and 50 µM RuBPY (tris(bipyridine)ruthenium(II) chloride). Media, based on DMEM D5030, contained 25mM glucose, 10% FBS, 1% PS, 1 mM pyruvate, 1% glutamax and 2 mM HEPES and 2 mM MES to provide a low but constant buffering power over the pH range studied. Prior to measurements, each well was sealed with 150 µM mineral oil to restrict O_2_ ingress. HPTS and RuBPY fluorescence were monitored for 17 h using a Cytation 5 device (BioTek, Agilent, Winooski, VT, USA). Excitation was provided by a monochromator, and fluorescence emission was detected sequentially at five wavelengths, which were optimized as described previously (32).

### Sulforhodamine B (SRB) and GFP growth assays

GFP-labelled cells (pLV-eGFP addgene plasmid # 36083) were transduced with lentiviral constructs. 48 h later cells were seeded at 2,000 cells/well on flat-bottom 96-well plates (Ibidi). For co-cultures, the ratio of the two cellular populations was varied. The following day, medium was replaced with bicarbonate-buffered media (D7777, Sigma), and incubated at 37°C with 5% CO_2_ for seven days; in some experiments, atmospheric O_2_ was reduced to 2% to produce hypoxia. For some experiments, incubations were terminated on days 2, 3, 4, 5, 6 and 7 to obtain a time course. Plates were analyzed for GFP fluorescence and SRB absorbance (Cytation 5). The GFP signal was measured in intact monolayers (excitation 490 nm, emission 520 nm). Next, wells were prepared for the SRB assay by fixing with PFA 4% for 30 min, washing with water and staining with 100 μL 0.057% SRB (in 1% acetic acid) for 30 min. Residual SRB was removed by washing with 1% acetic acid four times. Finally, 200 μL/well 10 mM Tris base was added to release SRB and measure absorbance (520 nm).

### Statistics

Data expressed as mean±SEM. Significance: *=P<0.05, **=P<0.01, ***=P<0.001. Gaussian mixture modelling was performed using GMMchi (https://github.com/jeffliu6068/GMMchi).

## ACKNOWLEDGEMENTS

The work was supported by the European Research Council, SURVIVE #723997. We thank Professor Alison Simmons and Dr Agne Antanaviciute for access to the single-cell RNA GSE datasets for colonic epithelium.

## AUTHOR CONTRIBUTIONS

SM, JM, AEH, AH performed experiments and data analysis, GA performed GMM analysis, WFB provided cell lines and advice on analysis, PS designed the research and wrote the paper.

## DATA AVAILABILITY STATEMENT

All data generated or analysed during this study are included in this published article and its supplementary information files.

---

**Supplementary Figure S1:**
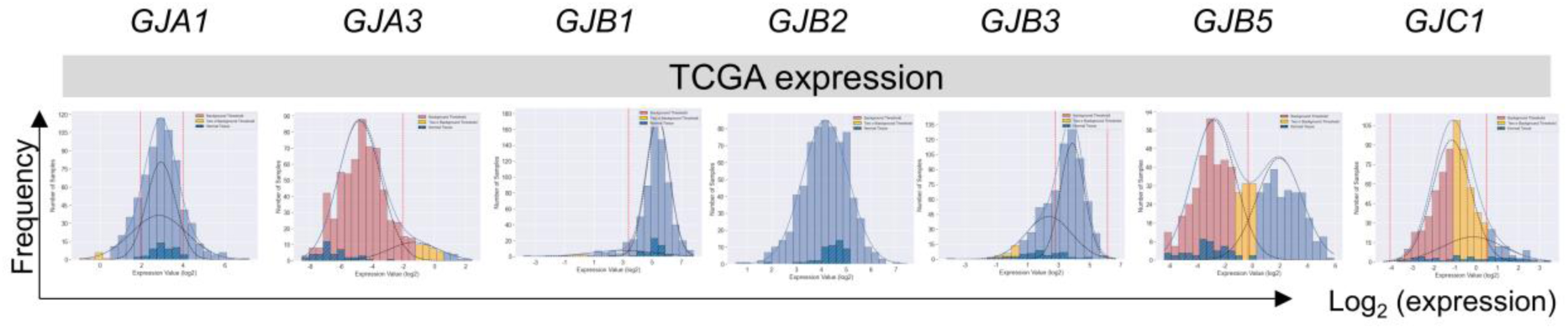
Application of GMMchi pipeline in purifying non-tumor expression from bulk tumor expression in the TCGA patient samples revealed that out of 689 TCGA CRC samples, 51 are paired normal samples and 637 tumor samples. Graphs show the expression distribution of the *GJA1*, *GJA3*, *GJB1*, *GJB2*, *GJB3*, *GJB5*, and *GJC1* combining paired tumor and normal samples.

**Supplementary Figure S2:**
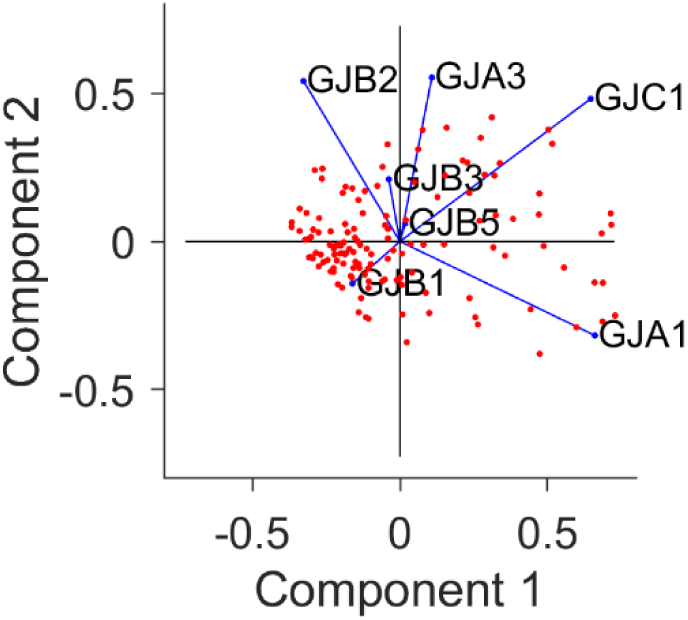
Principal component analysis (PCA) of microarray data showing weights for the seven Cx genes.

**Supplementary Figure S3:**
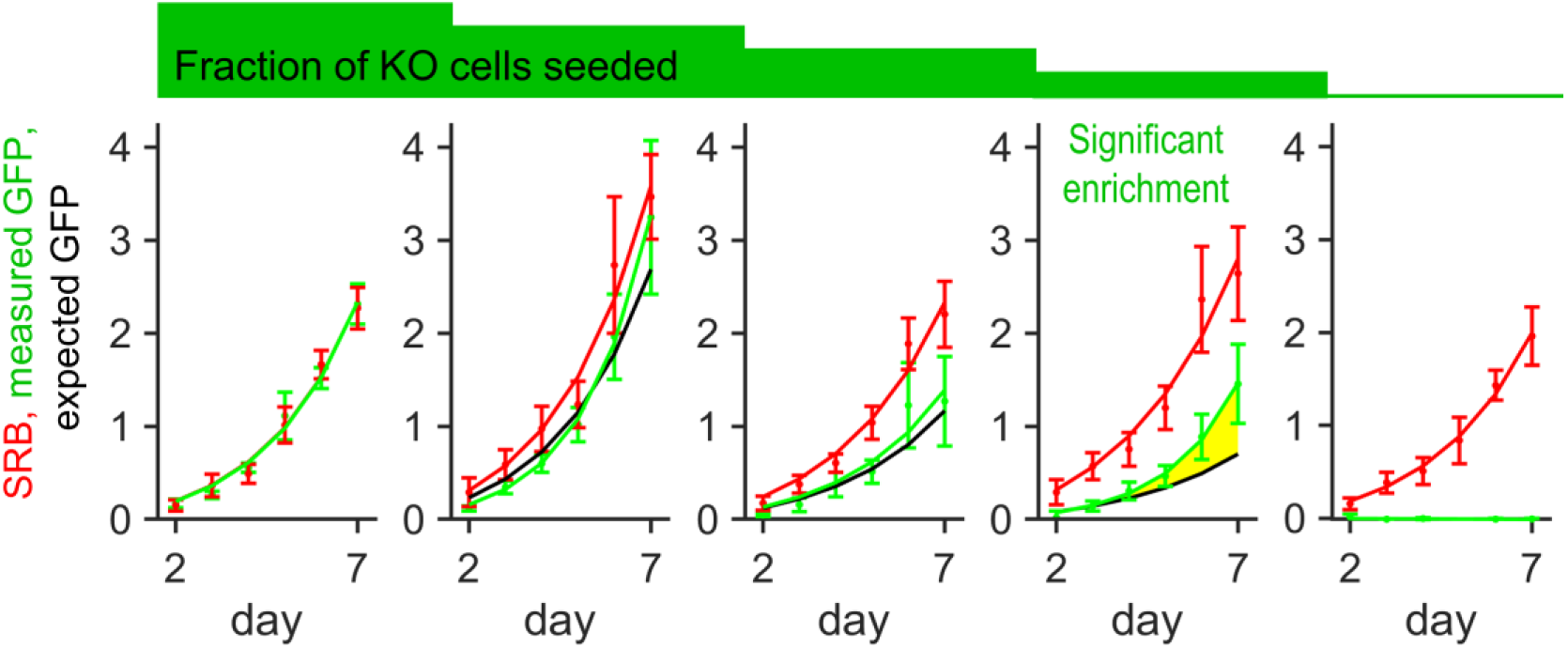
Growth curves for HCT116 cells grown as co-cultures of various ratios of WT and KO1 cells, quantified in terms of GFP fluorescence (KO compartment) and SRB absorbance (total biomass) over 7 days of culture, starting from a seeding density of 2,000 cells/well. Mean±SEM (N=5 per construct, with 3 technical repeats each). Significant enrichment indicates that the KO compartment expanded faster than expected from the SRB curve and seeding ratio (2,000:0, 1500:500, 1000:1000, 500:1500 and 0:2000), indicating that KO cells benefited from coupling onto WT neighbors. Statistical testing by two-way ANOVA; P value reported for difference between measured growth of KO cells (GFP) and prediction growth (SRB, scaled by KO:WT seeding ratio). Significance at P<0.05. Media were at pH 6.6.

**Supplementary Figure S4:**
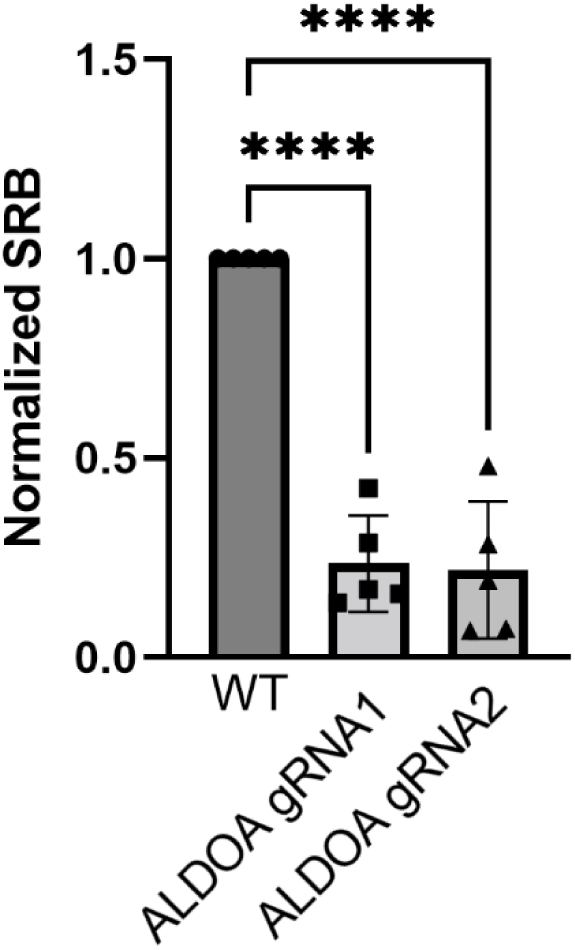
SRB absorbance after 6 days of culture of *ALDOA*-deficient cells infected with construct #1 or #2, normalized to time-matched WT cells. One-way ANOVA, pairwise testing.

**Supplementary Figure S5:**
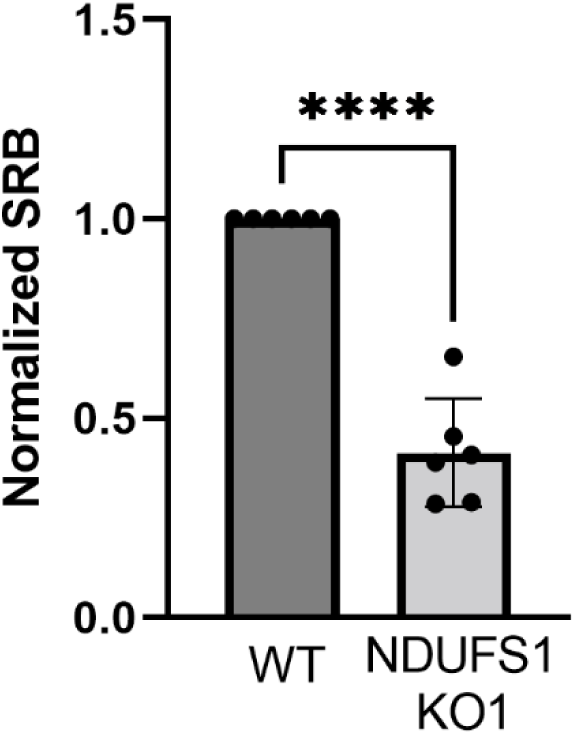
SRB absorbance after 6 days of culture of *NDUFS1*-deficient cells, normalized to time-matched WT cells. T-test.

